# Disproportionate presence of adenosine in mitochondrial and chloroplast DNA of *Chlamydomonas reinhardtii*

**DOI:** 10.1101/2020.08.27.270314

**Authors:** Waleed M. M. El-Sayed, Alli L. Gombolay, Penghao Xu, Taehwan Yang, Youngkyu Jeon, Sathya Balachander, Gary Newnam, Sijia Tao, Nicole E. Bowen, Raymond F. Schinazi, Baek Kim, Yongsheng Chen, Francesca Storici

## Abstract

Ribonucleoside monophosphates (rNMPs) represent the most common non-standard nucleotides found in the genomic DNA of cells. The distribution of rNMPs in DNA has been studied only in limited genomes, such as yeast nuclear and mitochondrial DNA, as well as human mitochondrial DNA. In this study, we used the ribose-seq protocol and the Ribose-Map bioinformatics toolkit to reveal the distribution of rNMPs incorporated into the whole genome of a photosynthetic unicellular green alga, *Chlamydomonas reinhardtii*. The study presents the discovery of a disproportionate incorporation of adenosine in the mitochondrial and chloroplast DNA, in contrast to the nuclear DNA, relative to the nucleotide content of these *C. reinhardtii* genomes. Our results demonstrate that the rNMP content in the DNA of the algal organelles reflects an elevated ATP level present in the algal cells. In addition, we reveal specific rNMP biases and patterns in the mitochondrial, chloroplast and nuclear DNA of *C. reinhardtii*.

## Introduction

The presence of ribose sugar in place of deoxyribose in DNA is a common DNA modification due to the abundant incorporation of ribonucleoside monophosphates (rNMPs), which are the units of RNA, by DNA polymerases (Nava et al., 2020; Williams et al., 2016). While it has been known for a long time that rNMPs are present in specific DNA sequences, such as mouse and human mitochondrial DNA (Grossman et al., 1973), at the mating type locus in the nuclear DNA of fission yeast (Vengrova and Dalgaard, 2006) and even in chloroplast DNA (Kolodner et al., 1975), only in the last decade has the ribose in DNA been defined as the most abundant alteration in the DNA of cells (Caldecott, 2014; Cavanaugh et al., 2010; Clausen et al., 2013; Gosavi et al., 2012; Kasiviswanathan and Copeland, 2011; Kennedy et al., 2012; Lemor et al., 2018; McDonald et al., 2012; Nick McElhinny et al., 2010; Potenski and Klein, 2014; Williams and Kunkel, 2014; Williams et al., 2016). Recent studies highlight the capacity of many DNA polymerases to incorporate rNMPs into DNA (Astatke et al., 1998; Bonnin et al., 1999; Brown and Suo, 2011; Cavanaugh et al., 2010; Gong et al., 2005; Kasiviswanathan and Copeland, 2011; Kennedy et al., 2012; McDonald et al., 2012; Nick McElhinny and Ramsden, 2003; Patel and Loeb, 2000). For example, *Escherichia coli* polymerase V (McDonald et al., 2012), the polymerase component of bacterial non-homologous end joining ligases (Zhu and Shuman, 2008), all replicative polymerases of budding yeast (Pol α, δ and ε) (Nick McElhinny et al., 2010), and the human replicative polymerase δ (Clausen et al., 2013) can insert rNMPs into DNA. Human DNA polymerases λ and μ can insert rNMPs with the same efficiency as deoxyribonucleoside monophosphates (dNMPs) (Gosavi et al., 2012; Moon et al., 2017). In addition, the reverse transcriptase of the human immunodeficiency virus inserts 1 rNMP per 146 dNMPs in the viral genome before integrating into human macrophage DNA (Kennedy et al., 2012). Although these data suggest that rNMPs in DNA are broadly present in nature, the primary attention to positions, patterns, and hotspots of rNMPs in DNA has been only in yeast genomic DNA (Balachander et al., 2020; Clausen et al., 2015; Daigaku et al., 2015; Jinks-Robertson and Klein, 2015; Koh et al., 2015; Reijns et al., 2015). Because ribonuclease (RNase) H2 is the major enzyme removing rNMPs from DNA (Sparks et al., 2012), the initial mapping of rNMPs in DNA to a single-nucleotide resolution was done in yeast strains with non-functional RNase H2 (Clausen et al., 2015; Daigaku et al., 2015; Jinks-Robertson and Klein, 2015; Koh et al., 2015; Reijns et al., 2015). Moreover, several reports have shown that RNase H2 is not active in yeast and human mitochondrial DNA (Balachander et al., 2020; Berglund et al., 2017; Wanrooij et al., 2017). This likely explains why, using a collection of *Saccharomyces cerevisiae* mutants with altered nucleotide pool composition, it was shown that variations in the dNTP pool can significantly affect rNMP frequency in mitochondrial DNA both in wild-type and RNase H2 defective cells, while the rNMP frequency was altered in nuclear DNA only in RNase H2-defective cells but not in wild-type cells (Wanrooij et al., 2017). Similarly, depletion of dNTP pools promoted rNMP incorporation in human mitochondrial DNA (Berglund et al., 2017). While nucleotide pool imbalances certainly play a major role in rNMP composition in genomic DNA, there are factors beyond variation in nucleotide pools that affect distribution and patterns of rNMP incorporation in DNA. Recently, by studying rNMP profiles in the genomes of three yeast species (*S. cerevisiae, Saccharomyces paradoxus* and *Schizosaccharomyces pombe*) and several different strains of these yeasts having either wild-type or mutant RNase H2, we showed low levels of rU in the mitochondrial DNA as well as in the nuclear DNA of RNase H2-defective cells of all three yeast species, as well as dominant rC and low rG in the nuclear DNA of wild-type RNase H2 cells of all three yeast species (Balachander et al., 2020). We further demonstrated non-random incorporation in both the yeast nuclear and mitochondrial DNA, uncovering the fact that the dNMP immediately upstream of the rNMP has a strong impact on the pattern of rNMP incorporation in mitochondrial as well as nuclear yeast DNA (Balachander et al., 2020).

The inclusion of rNMPs in DNA alters its stability, structure, plasticity, and ability to interact with proteins (Chiu et al., 2014; Klein, 2017). The presence of rNMPs in DNA may also regulate/modulate cellular functions. Thus, it is important to map rNMP sites in DNA and to characterize their features and rules of incorporation to understand the biological significance of rNMPs in DNA and determine whether these features and rules are conserved across different organisms or cell types. Our molecular and computational approaches, ribose-seq (Balachander et al., 2019) and Ribose-Map (Gombolay et al., 2019) allow for the efficient construction and analysis of genomic libraries derived from any DNA source containing rNMPs. Exploiting these techniques, we focused on the unicellular green alga of the species *Chlamydomonas reinhardtii*, which is broadly distributed worldwide in soil and freshwater. It is used to study photosynthesis and cell mobility (Sasso et al., 2018). *C. reinhardtii* is also used in production of biofuels (Sasso et al., 2018). We built ribose-seq libraries of rNMP incorporation from three independent cultures of *C. reinhardtii* cells grown in light. We found a strongly biased incorporation of rAMP in the mitochondrial and chloroplast DNA, and we characterized the overall genomic rNMP distribution in these algal cells.

## Results

### *C. reinhardtii* cells have a high ATP/dATP ratio

With the goal to study the composition and distribution of rNMPs in the DNA of *C. reinhardtii* cells, we first determined the concentration of NTPs and dNTPs in these algal cells. *C. reinhardtii* (strain CC-1690) cells were grown in light for 5-7 days to reach O.D. = 1.5-1.7 at 750 nm corresponding to 2×10^6^ cells/mL. Cells were lysed, and cell extracts were prepared for mass spectrometry analyses of the NTP (ATP, CTP, GTP and UTP) and dNTP (dATP, dCTP, dGTP and dTTP) pools. The mass spectrometry analyses revealed strong abundance of ATP in the *C. reinhardtii* cells (**Figure 1A**). While NTPs are generally more abundant than dNTPs in cells, with NTP concentrations being one to three orders of magnitude higher than those of dNTPs (Clausen et al., 2013; Ferraro et al., 2010), a key factor contributing to misincorporation of rNMPs in DNA is a variation in nucleotide pool concentrations resulting in an increased NTP/dNTP ratio for one or more nucleotides (Ferraro et al., 2010; Wanrooij et al., 2017). In *C. reinhardtii* cells, the ATP/dATP ratio was by far the largest of the NTPs/dNTPs ratios in these cells (**Figure 1B**), and significantly higher than all other NTP/dNTP ratios (CTP/dCTP, GTP/dGTP and UTP/dTTP, *P* = 0.0007, 0.004 and 0.0006, respectively). Compared to the previously recorded ATP/dATP ratios obtained for yeast cells, varying from ~100 to ~300 in *S. cerevisiae* and ~750 in *S. pombe* (Balachander et al., 2020; Clausen et al., 2013), and between ~130 and ~350 in human dividing cells, and ~1,450 in human non-dividing cells (Clausen et al., 2013; Ferraro et al., 2010; Traut, 1994), the ATP/dATP ratio that we measured in *C. reinhardtii* cells is the highest, ~1,800 (**Figure 1B**).

**Figure 1.**
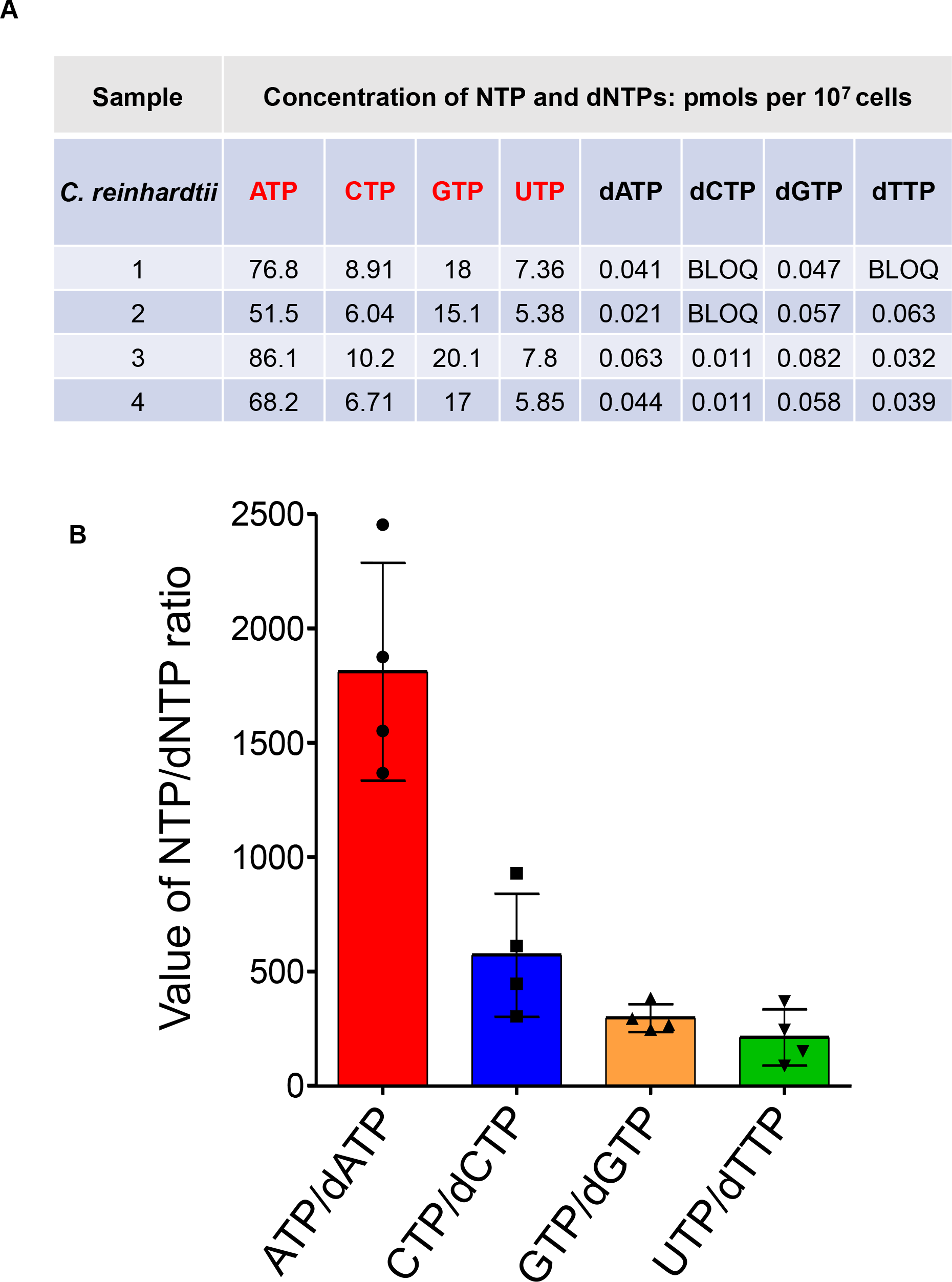
Measurements of NTP and dNTP levels in *C. reinhardtii* cells show high ATP concentration. (**A**) The levels of NTPs and dNTPs extracted from known numbers of algal cells of *C. reinhardtii* strain CC-1690 were determined by LC-MS/MS methods (see Methods). BLOQ: below limit of quantification. The cellular level of each nucleotide was normalized for 10^7^ cells. (**B**) The NTP and dNTP levels determined in A) were used to calculate the NTP/dNTP ratios. Shown are mean and standard deviation of four repeats for each sample. Ratios for dCTP and dTTP levels below the limit of quantification were calculated by using the limit value of quantification for dCTP or dTTP (0.02 pmole) relative to the number of cells used in the LC-MS/MS method. The ratio of ATP/dATP was clearly significantly higher than all other NTP/dNTP (CTP/dCTP, GTP/dGTP and UTP/dTTP) ratios, *P* = 0.0007, 0.004 and 0.0006, respectively.

### Mitochondrial and chloroplast DNA of *C. reinhardtii* have overriding incorporation of rA

To determine the pattern of rNMPs in the mitochondrial, chloroplast and nuclear genome of *C. reinhardtii* cells, we extracted the whole genomic DNA from three independent cultures of these cells grown in light (see Materials and Methods). From these three DNA extracts, we constructed three genomic libraries using the ribose-seq approach (see Materials and Methods and **Table S1**): FS121, FS231 and FS232 (**Table S2**). Each of these ribose-seq libraries was sequenced and then segmented into a mitochondrial, chloroplast and nuclear library using the Ribose-Map computational toolkit after alignment of the sequencing reads to the reference genome sequence of *C. reinhardtii*. We obtained the mitochondrial and chloroplast sequences from NCBI (https://www.ncbi.nlm.nih.gov/genome/147), and the 5.5 nuclear sequence from JGI (https://phytozome.jgi.doe.gov/pz/portal.html#!info?alias=Org_Creinhardtii). The percentage of rA, rC, rG and rU varies strikingly between ribose-seq libraries of the organelles and nuclear DNA. Remarkably, both the mitochondrial and chloroplast DNA display a noticeable preference for rA with an average of 89% rA in mitochondrial and 77% rA in chloroplast DNA (**Figure 2** and **Table S2**). As shown by our analysis of the rNMP content in mitochondrial and chloroplast DNA of *C. reinhardtii* cells, rA is significantly and disproportionally incorporated relative to rC, rG, rU and relative to the nucleotide content of these genomes (**Figures 3A-D**, **4A,B** and **Figure 5A**, and **Table S3A,B**). The biased incorporation of rA in these organelles likely reflects the high ratio of ATP/dATP in the algal cells (**Figure 1B**). Interestingly, while rC, rG and rU are all similarly infrequent in the mitochondrial DNA, rU is distinctly the least abundant rNMP in the chloroplast DNA, even if rC and rG are much less frequent than rA, as evidenced via nucleotide frequency and heatmap analyses (**Figures 3A-D**, **4A,B** and **Figure 5A**, and **Table S3A,B**).

**Figure 2.**
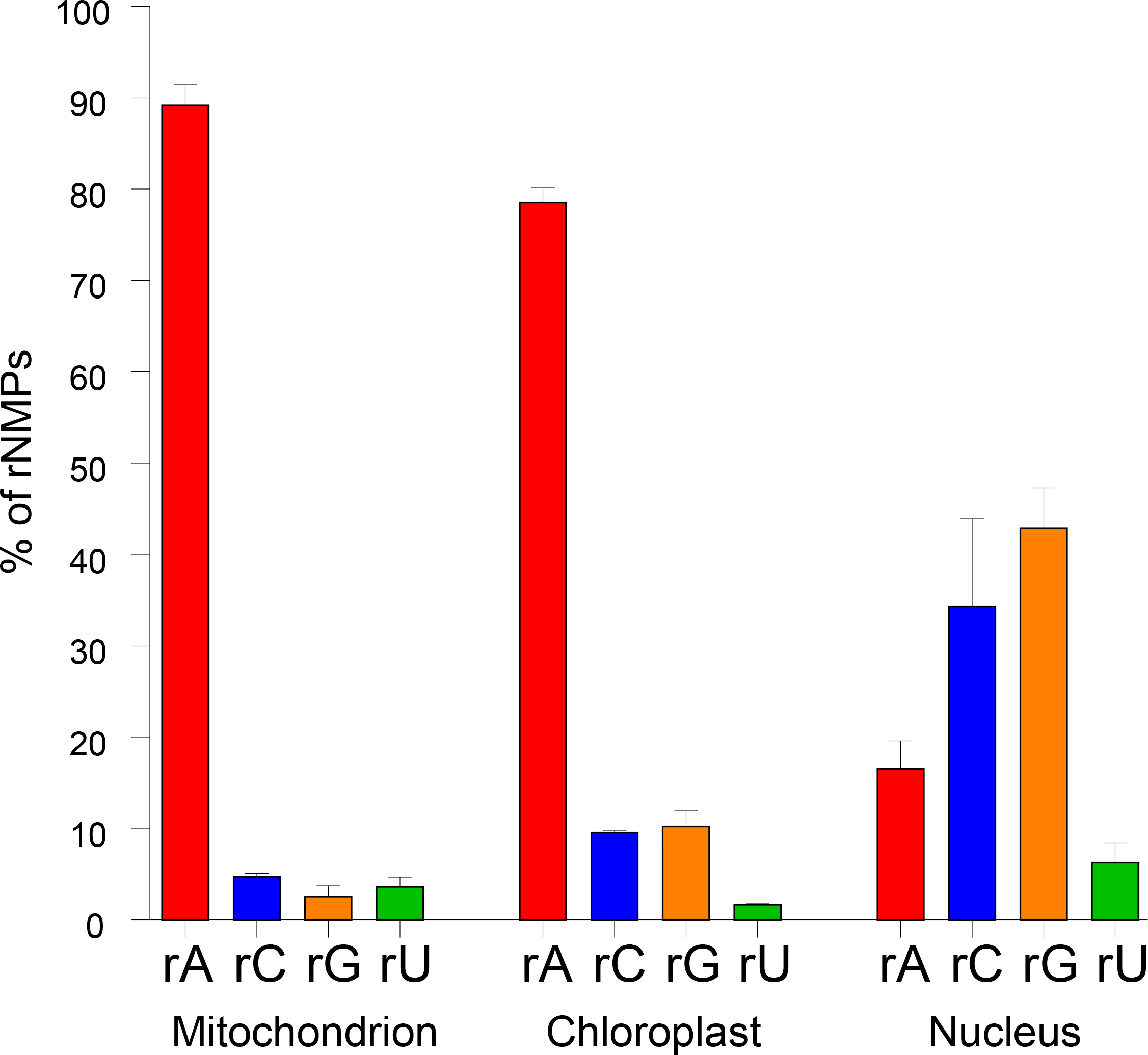
High frequency of rA in mitochondrial and chloroplast but not nuclear genome of *C. reinhardtii* cells. Bar graph with percentage of rA, rC, rG and rU found in mitochondrial, chloroplast and nuclear DNA of *C. reinhardtii* cells. Mean and standard deviation from three different mitochondrial, chloroplast or nuclear ribose-seq libraries are shown.

**Figure 3.**
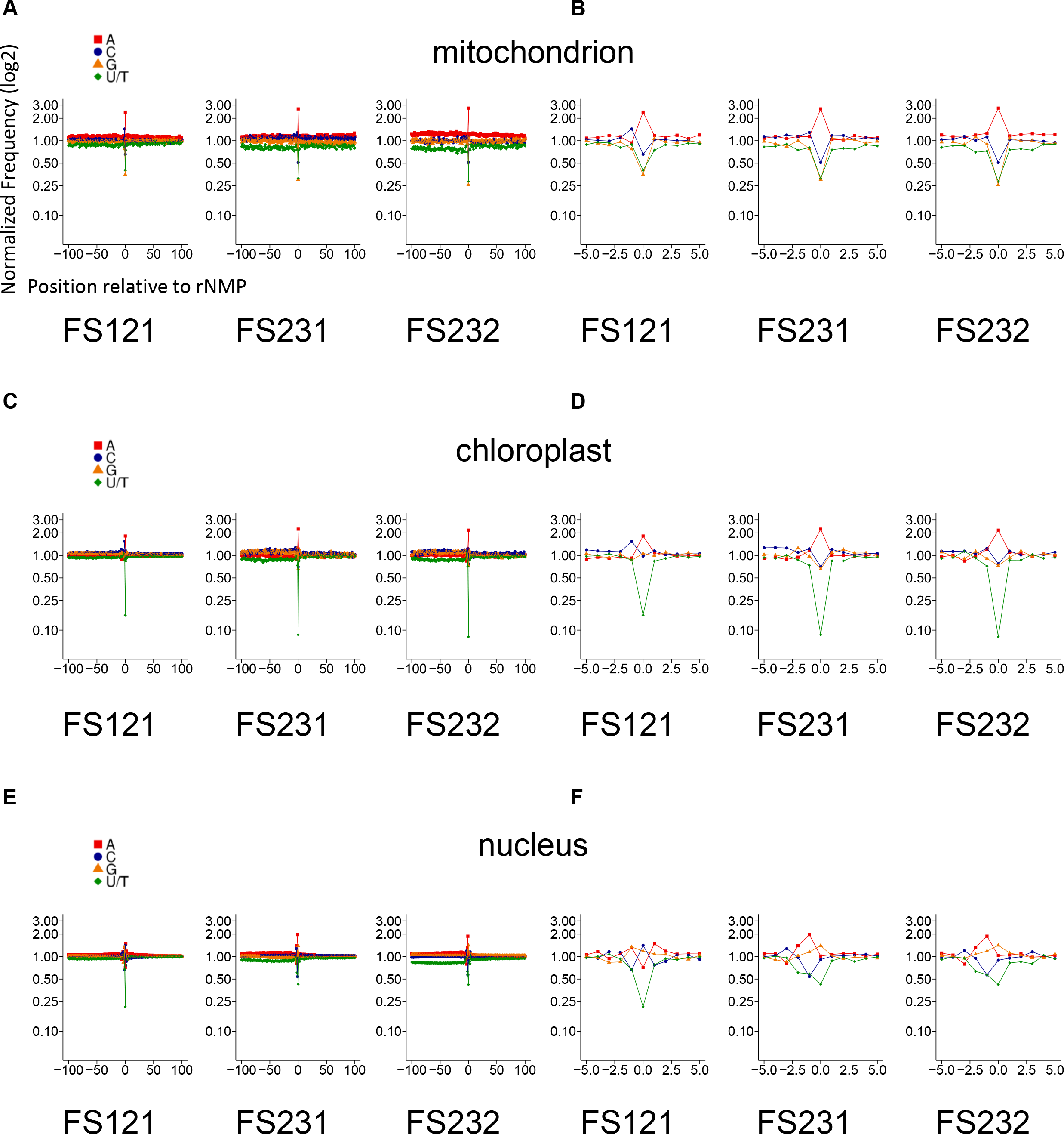
Identity and sequence context of rNMPs in mitochondrial, chloroplast and nuclear DNA of *C. reinhardtii* cells. Zoomed-out (**A**, **C** and **E**) and zoomed-in (**B**, **D** and **F**) plots of normalized nucleotide frequencies relative to mapped positions of sequences from mitochondrial (**A** and **B**), chloroplast (**C** and **D**) and nuclear (**E** and **F**) ribose-seq libraries. Position 0 is the rNMP, - and + positions are upstream and downstream dNMPs, respectively, normalized to the A, C, G and T content in the genome of *C. reinhardtii*. The y-axis shows the frequency of each type of nucleotide present in the ribose-seq data normalized to the frequency of the corresponding nucleotide present in the reference genome of the indicted cell compartment of *C. reinhardtii*. Red square, A; blue circle, C; orange triangle, G; and green rhombus, U. Red square, A; blue circle, C; orange triangle, G; and green rhombus, U.

**Figure 4.**
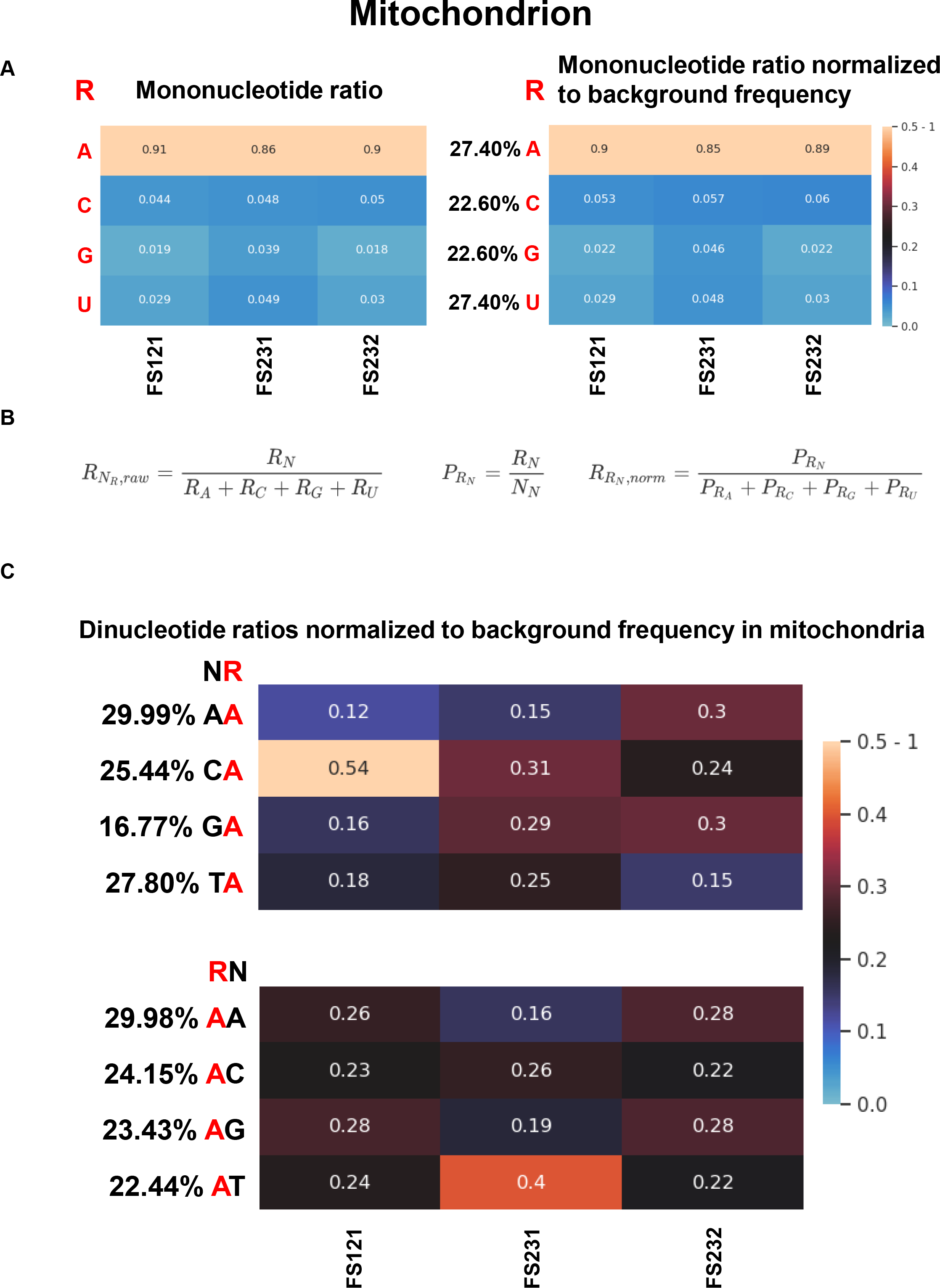
rA is by large the most abundantly incorporated rNMP in *C. reinhardtii* mitochondrial DNA. (**A**) Heatmap analyses with (left) ratio of each type of rNMP (rA, rC, rG and rU), and (right) ratio of each type of rNMP normalized to the nucleotide frequencies of the *C. reinhardtii* mitochondrial reference genome for the mitochondrial ribose-seq libraries of this study. The corresponding formulas used are shown in B) and explained in the Materials and Methods. Each column of the heatmap shows results of a specific ribose-seq library. Each library name is indicated underneath each column of the heatmap. Each row shows results obtained for an rNMP (R in red) of base A, C, G or U for each library. The actual percentage of A, C, G and T bases present in mitochondrial DNA of *C. reinhardtii* are shown to the left of the heatmap with normalized data. The bar to the right shows how different ratio values are represented as different colors. Black corresponds to 0.25. (**B**) Formulas used to calculate the ratio and the normalized ratio values of the mononucleotide heatmaps for mitochondrial in A) above, chloroplast (**Figure 5A**), and nuclear (**Figure 6A**) DNA. (**C**) Heatmap analyses with normalized frequency of mitochondrial NR (top) and RN (bottom) dinucleotides containing rA with the upstream (top), or downstream (bottom) deoxyribonucleotide with base A, C, G or T for the mitochondrial ribose-seq libraries of this study. The formulas used to calculate these normalized frequencies are shown and explained in the Materials and Methods. Each column of the heatmap shows results of a specific ribose-seq library. Each library name is indicated underneath each column of the heatmap. Each row shows results obtained for a dinucleotide NR or RN (R in red) of fixed rNMP base A for each library. The actual % of dinucleotides of fixed base A for the indicated base combinations (AA, CA, GA and TA, top; and AA, AC, AG and AT, bottom) present in mitochondrial DNA of *C. reinhardtii* are shown to the left of the corresponding heatmaps. The observed % of dinucleotides with rA were divided by the actual % of each dinucleotide with fixed base A in mitochondrial DNA of *C. reinhardtii*. The bar to the right shows how different frequency values are represented as different colors. Black corresponds to 0.25. Significance of comparisons for data in this Figure are shown in **Table S3A**.

**Figure 5.**
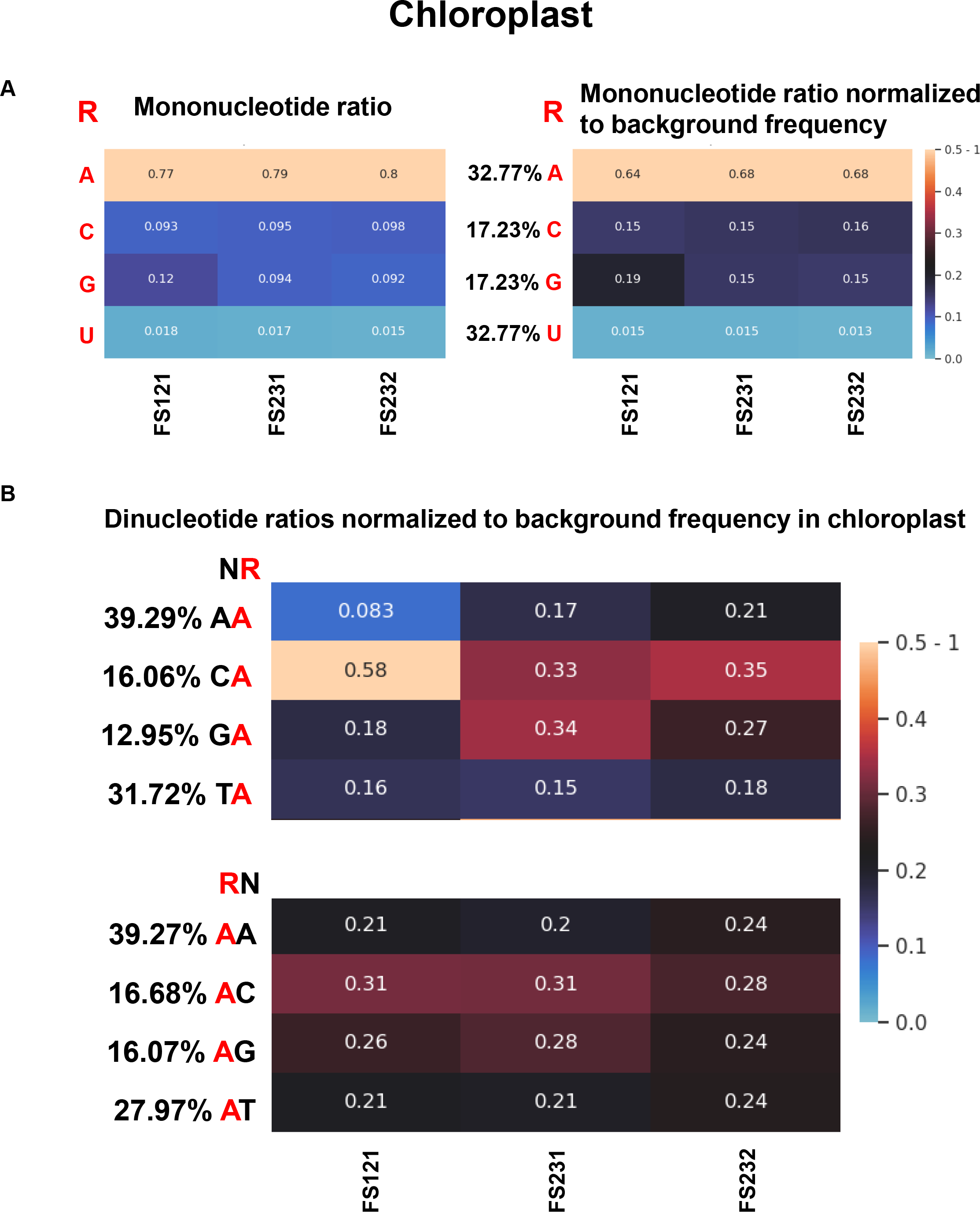
rA is dominant and not randomly distributed in *C. reinhardtii* chloroplast DNA. (**A**) Heatmap analyses with (left) ratio of each type of rNMP (rA, rC, rG and rU), and (right) ratio of each type of rNMP normalized to the nucleotide frequencies of the *C. reinhardtii* chloroplast reference genome for the chloroplast ribose-seq libraries of this study. The corresponding formulas used are shown in **Figure 4B**, and explained in the Materials and Methods. Each column of the heatmap shows results of a specific ribose-seq library. Each library name is indicated underneath each column of the heatmap. Each row shows results obtained for an rNMP (R in red) of base A, C, G or U for each library. The actual percentage of A, C, G and T bases present in chloroplast DNA of *C. reinhardtii* are shown to the left of the heatmap with normalized data. The bar to the right shows how different ratio values are represented as different colors. Black corresponds to 0.25. (**B**) Heatmap analyses with normalized frequency of chloroplast NR (top) and RN (bottom) dinucleotides containing rA with the upstream (top), or downstream (bottom) deoxyribonucleotide with base A, C, G or T for the chloroplast ribose-seq libraries of this study. The formulas used to calculate these normalized frequencies are shown and explained in the Materials and Methods. Each column of the heatmap shows results of a specific ribose-seq library. Each library name is indicated underneath each column of the heatmap. Each row shows results obtained for a dinucleotide NR or RN (R in red) of fixed rNMP base A for each library. The actual % of dinucleotides of fixed base A for the indicated base combinations (AA, CA, GA and TA, top; and AA, AC, AG and AT, bottom) present in chloroplast DNA of *C. reinhardtii* are shown to the left of the corresponding heatmaps. The observed % of dinucleotides with rA were divided by the actual % of each dinucleotide with fixed base A in chloroplast DNA of *C. reinhardtii*. The bar to the right shows how different frequency values are represented as different colors. Black corresponds to 0.25. Significance of comparisons for data in this Figure are shown in **Table S3B**.

**Figure 6.**
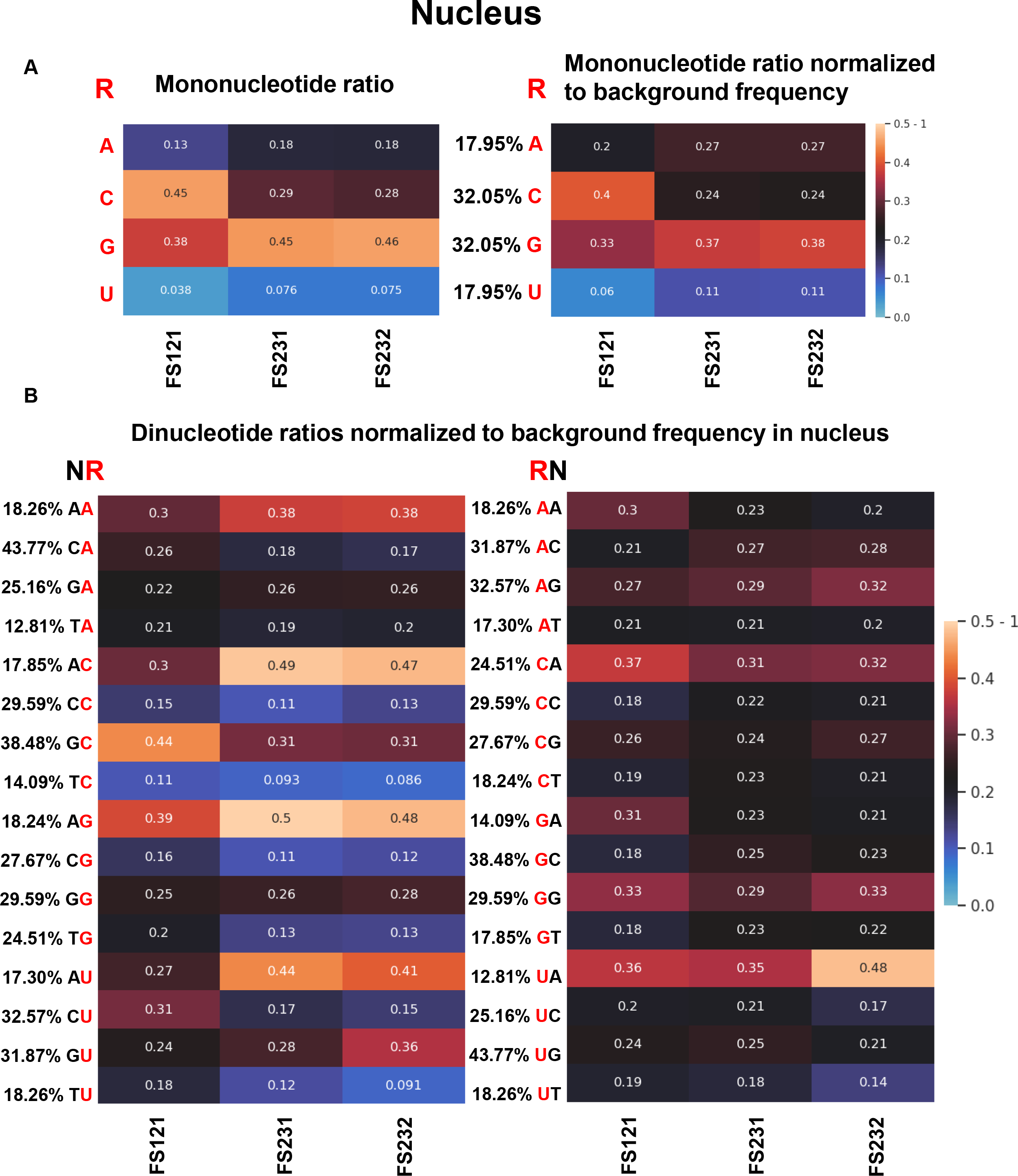
rC and rG are dominant and not randomly distributed in nuclear DNA of *C. reinhardtii* cells. (**A**) Heatmap analyses with (left) ratio of each type of rNMP (rA, rC, rG and rU), and (right) ratio of each type of rNMP normalized to the nucleotide frequencies of the *C. reinhardtii* nuclear reference genome for the nuclear ribose-seq libraries of this study. The corresponding formulas used are shown in **Figure 4B**, and explained in the Materials and Methods. Each column of the heatmap shows results of a specific ribose-seq library. Each library name is indicated underneath each column of the heatmap. Each row shows results obtained for an rNMP (R in red) of base A, C, G or U for each library. The actual percentage of A, C, G and T bases present in nuclear DNA of *C. reinhardtii* are shown to the left of the heatmap with normalized data. The bar to the right shows how different ratio values are represented as different colors. Black corresponds to 0.25. (**B**) Heatmap analyses with normalized frequency of nuclear NR (top) and RN (bottom) dinucleotides with rA, rC, rG and rU with the upstream (left), or downstream (right) deoxyribonucleotide with base A, C, G or T for the nuclear ribose-seq libraries of this study. The formulas used to calculate these normalized frequencies are shown and explained in the Materials and Methods. Each column of the heatmap shows results of a specific ribose-seq library. Each library name is indicated underneath each column of the heatmap. Each row shows results obtained for a dinucleotide NR or RN (R in red) of fixed base A, C, G or T for the indicated base combinations for each library. The actual % of dinucleotides of fixed base A, C, G or T present in nuclear DNA of *C. reinhardtii* are shown to the left of the corresponding heatmaps. The observed % of dinucleotides with rNMPs with base A, C, G or U were divided by the actual % of each dinucleotide with fixed base A, C, G or T in nuclear DNA of *C. reinhardtii*. The bar to the right shows how different frequency values are represented as different colors. Black corresponds to 0.25. Significance of comparisons for data in this Figure are shown in **Table S3C**.

### Nuclear DNA of *C. reinhardtii* has higher level of rG and rC and lower level of rA and rU

The distribution of rNMPs in nuclear DNA showed more abundant rG and rC, followed by rA (**Figures 2, 3E,F** and **6A**, and **Table S2**). rU was consistently the least abundant rNMP in the nuclear DNA of the three libraries, and on average a factor of ~2.8 and up to a factor of 4.7 below the nuclear-dT content (**Figures 3E,F** and **6A**, and **Tables S2 and S3C**). Normalization of single rNMP frequencies to the nucleotide base content of *C. reinhardtii* nuclear DNA revealed non-random rNMP incorporation with a marked preference for rG and/or rC in all of the nuclear libraries over rA, and especially over rU (**Figure 6A** and **Table S3C**). These nuclear data do not reflect the nucleotide pool composition of *C. reinhardtii* cells, because, as shown above, among the measured ratios of NTPs/dNTPs in the algal cells, the ATP/dATP is the highest one (**Figure 1B**). These results suggest possible active removal of rA from nuclear DNA, but not from the mitochondrial and chloroplast genomes. Therefore, we examined whether the *C. reinhardtii* cells of strain CC-1690 displayed RNase H2 activity or other similar activity on a DNA substrate containing an embedded rNMP. We prepared protein extracts (**Figure S1A**) and examined the activity of the extracts to cleave a substrate with one rG or one rA (see Materials and Methods). Cleavage of these substrates by *E. coli* RNase HII was used as a positive control. The results show that *C. reinhardtii* protein extract is not active on an rG in DNA because the dsDNA substrate was not cleaved at the rNMP position (**Figure S1B**). Similar results were obtained using the DNA substrate containing rA (**Figure S1B**). No active cleavage of rNMP-containing substrates from the *C. reinhardtii* protein extract was detected. It is possible that the RNase H2 activity in *C. reinhardtii* cells is weak. At the same time, we cannot exclude the possibility that the protein extract is either missing some cofactor necessary for the RNase H2 function of *C. reinhardtii*, or it may contain some inhibitory factor for RNase H2, preventing RNase H2 from cutting the rNMP-containing substrates. When we treated the substrate containing the rG with *E. coli* RNase HII both in the presence and the absence of the algal protein extract, we noticed a modest decrease of RNase HII cleavage activity in the presence of the algal protein extract, although cleavage at the rG was still efficient (**Figure S1C**).

### rA found in mitochondrial and chloroplast DNA, and rC, rG and rA found in nuclear DNA are not randomly incorporated

We then studied whether rA is randomly incorporated in the mitochondrial and chloroplast DNA of the three *C. reinhardtii* libraries. If, for example rA is randomly incorporated in *C. reinhardtii* mitochondrial DNA, the frequency by which the dNMP with base A, C, G or T is found at position −1 and +1 relative to rA should reflect the frequency of the dinucleotides AA, CA, GA, TA, AC, AG and AT obtained from the sequence of *C. reinhardtii* mitochondrial DNA. The heatmap dinucleotide analysis revealed that the frequency of the dNMPs immediately upstream (at position −1) of the rAMP, and in part the dNMP at position +1 and −3 from the rAMP in mitochondrial DNA, deviates from the expected values but varies among the three mitochondrial libraries (**Figure 4C**, **Figure S2A**, and **Table S3A**). No particular bias was found for the nucleotide upstream or downstream of the rAMP at positions −2, +2, +3, −4,+4, −5, +5, −6, +6, −100 and +100 in the mitochondrial DNA (**Figure S2A**). In fact, with the increased gap between dinucleotide pairs (from −2, +2 to −6, +6, and −100 and +100), the randomness is weakened, as evidenced by an increased *P*-value from −1, +1 to −100 and +100 (**Table S3A**). For the chloroplast DNA, the rAMP was preferentially found downstream of dC and more rarely downstream of dA and dT (**Figure 5B** and **Table S3B**). No particular bias was found for the nucleotide upstream or downstream of the rAMP at positions +1, −2, +2, −3, +3, −4,+4, −5, +5, −6, +6, −100 and +100 in the chloroplast DNA (**Figure S2B** and **Table S3B**). Due to the low abundance of rC, rG and rU in the mitochondrial and chloroplast DNA, we did not analyze the surrounding dNMPs of these rNMPs.

For nuclear DNA, we found that both rC and rG are preferentially preceded by dA at position −1. The observed count of NR dinucleotides with an rC or rG (dArC, dCrC, dGrC and dTrC; dArG, dCrG, dGrG and dTrG) was significantly different from the expected count calculated for the background frequency of the corresponding dinucleotide pair with the same number of total rCMPs or rGMPs (**Table S3C**). The observed count of the dinucleotide dArC was above the expected count for this pair **(Figure 6B** and **Table S3C**. Similarly, the observed count of dArG was above the expected count for this pair **(Figure 6B** and **Table S3C**). For rA, we found that the dArA count was above the expected count for this dinucleotide in all libraries, while for rU the pattern was less clear, possibly due to the fact that rU is the least abundant rNMP found in the nuclear DNA of *C. reinhardtii* cells and more data would be needed to obtain an accurate spectrum of incorporation (**Figure 6B** and **Table S3C**). A much less prominent difference was found among pair combinations for the dNMPs at position +1 (**Figure 6B** and **Table S3C**). We also examined the dNMPs at positions −2, +2, −3, +3, −4,+4, −5, +5, −6, +6, −100 and +100. The heatmaps progressively became uniformly darker from −2, +2 to −100, +100, while no particular pattern emerging (**Figure S3** and **Table S3C**). Overall, these results, which were also conserved among all three nuclear libraries, highlight the fact that the dNMPs immediately upstream of the rNMP at position −1 have the most impact on the incorporation of a specific rNMP type in a given genomic position of the algal nuclear DNA. Moreover, these findings demonstrate that rNMPs are not randomly incorporated in the nuclear genome of *C. reinhardtii* cells.

## Discussion

We report a genome-wide analysis of rNMP sites in a photosynthetic organism, the unicellular freshwater green alga *C. reinhardtii*. We found rNMPs embedded in all genomes of the alga: mitochondrial, chloroplast and nuclear. We revealed strikingly biased rA incorporation in the mitochondrial and chloroplast DNA, but not in the nuclear DNA, in which instead rG and rC are dominant over rA and rU. The disproportionate presence of rA embedded in the mitochondrial and chloroplast DNA reflects the remarkably high ATP/dATP ratio of the cells. These findings support the lack of RNase H2 activity in mitochondrial DNA and provide new evidence for the absence of RNase H2 activity on rNMPs embedded in chloroplast DNA. The inability to cleave and initiate removal of rNMPs embedded in mitochondrial and chloroplast DNA likely allows abundant incorporation of rNMPs in these genomes, particularly rA, significantly and overwhelmingly above the frequency expected based on the dA-content of the mitochondrial and chloroplast genomes. Work in yeast cells has provided supportive evidence for a frequent exchange of nuclear and mitochondrial dNTP pools (Wanrooij et al., 2017). Thus, we would expect rA to also be highly incorporated in the nuclear genome of *C. reinhardtii*. Because rA was not found to be the most frequently incorporated rNMP in the nuclear DNA of *C. reinhardtii*, but rather rG and rC were more frequently detected, our findings suggest that RNase H2 might have a high workload in removing rA from the nuclear genome of a photosynthetic organism, such as *C. reinhardtii*. Nevertheless, we were unable to detect an RNase H2 or any other cleavage function from protein extracts of the algal cells to support active removal of rA from nuclear DNA. It will be valuable to work with a purified RNase H2 enzyme from *C. reinhardtii* cells to characterize its activity on rNMPs embedded in DNA. There is also the possibility that the nuclear DNA polymerases of *C. reinhardtii* have a much stronger discrimination capacity for ATP vs. dATP than the DNA polymerases of the mitochondria and chloroplast of the alga. Further studies are needed to understand how rA is specifically excluded from the nuclear DNA of *C. reinhardtii* cells. Moreover, it would also be interesting to investigate the relation between rNMP incorporation in the mitochondrial and chloroplast DNA of the alga and the process of photosynthesis.

If we compare the rNMP content in the mitochondrial DNA of *C. reinhardtii* with that of *S. cerevisiae, S. paradoxus* and *S. pombe*, we find that rU is the least incorporated rNMP not only in mitochondrial and chloroplast DNA of the alga, as in the yeast mitochondria, but also in nuclear DNA of the alga. Differently, in yeast nuclear DNA, rU is the least frequent rNMP only in RNase H2-defective cells (Balachander et al., 2020). In part, the low level of rU incorporation could reflect the relatively low UTP/dTTP ratio in these eukaryotic cells. Activity of topoisomerase I on sequences with rU (Cho and Jinks-Robertson, 2018; Klein, 2017) and some proofreading activity for nuclear DNA polymerase on rU (Koh et al., 2015) could contribute to the general rare presence of rU in these genomes. Moreover, in each of the three yeast species, rA is incorporated significantly below the expected values for the corresponding genomes. Instead, rC or rG are the predominant rNMPs (Balachander et al., 2020). Our results show that the high ATP/dATP ratio in *C. reinhardtii* cells reflects the composition of rNMPs in the mitochondrial and chloroplast DNA of *C. reinhardtii*. Interestingly, rA incorporation was found mainly downstream of dC and/or dG in chloroplast DNA. However, no conserved pattern around rA was found in the mitochondrial DNA of the three ribose-seq libraries of *C. reinhardtii*. At the same time, rA did display a preference of incorporation downstream of specific dNMPs in the mitochondrial DNA of the three libraries, but such preference was not preserved in all three mitochondrial libraries. It is possible that small variations in growth conditions may affect the incorporation pattern of rA in the algal mitochondrial DNA. While rU is consistently the least represented rNMP in the nuclear, mitochondrial and chloroplast DNA of *C. reinhardtii* cells, the mitochondrial DNA also showed very low rG and rC content. Thus, despite the pervasive rA incorporation is a common feature in the mitochondrial and chloroplast DNA of *C. reinhardtii*, the overall patterns of rNMP incorporation in these two algal organelles are not identical, suggesting different rNMP incorporation mechanisms in these two organelles or variations in local nucleotide pools.

Another conserved feature between yeast and *C. reinhardtii* rNMP patterns, particularly for the nuclear DNA of the alga, is that the dNMP immediately upstream from the rNMP is the one that has the largest impact on the distribution of rNMPs. In fact, with the increased gap between dinucleotide pairs with an rNMP (from −1, +1 to −100, +100), the randomness is weakened not only for nuclear rNMPs, but also for the mitochondrial and chloroplast rNMPs, as shown by increased *P*-value from −1, +1 dinucleotides to −100 and +100 dinucleotides (**Table S3A-C**). Similarly to what we found for rNMPs in the yeast genome (Balachander et al., 2020), we believe that this lack of unpredictability of rNMPs in the algal genome supports an accommodation mechanism by DNA polymerases that facilitates incorporation of rNMPs following specific dNMPs.

In conclusion, via mapping and genome-wide analysis of ribose-seq libraries of the unicellular green alga *C. reinhardtii*, we have revealed a unique distribution of rNMPs embedded in the DNA of the algal organelles compared to the nuclear DNA of the same cells. It will be interesting to characterize how such disproportionate presence of rA in the genome of the organelles of this photosynthetic organism impacts the DNA metabolic functions of these genomes during their day and night cycles.

## Acknowledgments

We thank the Molecular Evolution Core with A. Bryksin at the Parker H. Petit Institute for Bioengineering and Bioscience of the Georgia Institute of Technology for high throughput sequencing, M.D. Herron and C. Lindsey for supplemental aliquots of *C. reinhardtii* CC-1690 cells, K. Mukherjee and D. Kundnani for critically reading the manuscript, and all members of the Storici laboratory for assistance and feedback on this study. We acknowledge funding from the Egyptian Cultural Affairs & Missions Sector, Cairo, Egypt (to W.M.M.E.), AI136581 (to B.K.), AI150451 (to B.K.), MH116695 (to. R.F.S), the National Institutes of Health (R01ES026243 to F.S.) and the Howard Hughes Medical Institute (Faculty Scholar grant 55108574 to F.S.) for supporting this work.

## Author Contributions

Conceptualization, F.S. and W.M.M.E.; Methodology, W.M.M.E., A.L.G., P.X., T.Y., Y.J., S.B., G.N., S.T., N.E.B, Y.C. and F.S.; Investigation, W.M.M.E., A.L.G., P.X., T.Y., Y.J., S.B. and F.S.; Writing – Original Draft F.S.; Writing – Review & Editing W.M.M.E., A.L.G., P.X., T.Y., Y.J., S.B. and G.N.; Funding acquisition, R.F.S., B.K. and F.S. Resources, R.F.S., B.K., Y.C. and F.S. All authors commented on and approved the manuscript.

## STAR★Methods

## RESOURCE AVAILABILITY

### Lead Contact

Further information and requests for resources and reagents should be directed to and will be fulfilled by the Lead Contact, Francesca Storici.

### Materials Availability

All unique/stable reagents generated in this study are available from the Lead Contact.

### Data and Code Availability

The authors declare that the data supporting the findings of this study are available within the paper and its supplementary information files. The Ribose-Map bioinformatics toolkit is available for download at GitHub (https://github.com/agombolay/ribose-map). See also(Gombolay et al., 2019). Custom Python3 scripts for background subtraction is available for download at GitHub under GNU GPL v3.0 license (https://github.com/xph9876/ArtificialRiboseDetection). The custom Python3 scripts for heatmaps is available for download at GitHub under GNU GPL v3.0 license (https://github.com/xph9876/RibosePreferenceAnalysis)

Table S2 contains raw data. Bar graphs representing the percentage of each type of rNMP were made using GraphPad Prism 5 (GraphPad Software). The nucleotide sequence context plots were created using custom R scripts. All data generated in this study are available from the Lead Contact.

## EXPERIMENTAL MODEL AND SUBJECT DETAILS

### Algal model growing culture and conditions

The algae *Chlamydomonas reinhardtii* CC-1690 (wild type, mt+ strain) was obtained from the Chlamydomonas Resource Center at the University of Minnesota. Briefly, *C. reinhardtii* cells were grown in tris-acetate-phosphate medium (TAP) in a glass tube (5.1 cm width and 64 cm height) (Gorman and Levine, 1965). The culture was given continuous illumination from eleven 40-Watt 1.2 m long cool white fluorescent light bulbs with a light path of approximately 7.6 cm, and sparged with air at ~1 L/min at room temperature (approximately 25 °C). The Cole-Parmer Model 300 pH/ORP meter (Cole-Parmer Instrument Co., Vernon Hills, IL) was used with a temperature controller, i.e., the Wahl Model C962 1/8-DIN Controller, which were connected to a solenoid; CO_2_ was dosed periodically to maintain the pH at 8.0±0.2. *C. reinhardtii* cells were grown for 5-7 days to reach OD = 1.5-1.7 at 750 nm corresponding to 2×10^6^ cells/mL. Small cultures for preparation of nucleotide or protein extracts were grown in TAP medium in the light for 3-4 days.

## METHOD DETAILS

### dNTP and rNTP measurements

*C. reinhardtii* cell lysates were prepared to extract the NTPs and rNTPs as described in (Diamond et al., 2004) with some modifications. Briefly, *C. reinhardtii* cells were grown in 20 mL of TAP medium in the light for 3-4 days. Cell were harvested, washed with 1 x PBS and resuspended in 65% ice-cold methanol and mixed by pipetting. Cells were then vortexed for 2 min and then heated to 95 °C for 3 min then placed on ice for 1 min. The cells were spun at 14,000 x g for 3 min. Aliquots were lyophylized and stored at −80 °C. We quantified the intracellular dNTPs and rNTPs using an ion pair chromatography-tandem mass spectrometry method (Fromentin et al., 2010) with some modifications. Chromatographic separation and detection were conducted on a Vanquish Flex system (Thermo Scientific, Waltham, MA) coupled with a TSQ Quantiva triple quadrupole mass spectrometer (Thermo Scientific, Waltham, MA). Analytes were separated using a Kinetex EVO-C18 column (100 × 2.1 mm, 2.6 μm) (Phenomenex, Torrance, CA) at a flow rate of 250 μL/min. The mobile phase A consisted of 2 mM of ammonium phosphate monobasic and 3 mM of hexylamine in water and the mobile phase B consisted of acetonitrile. The LC gradient increased from 10% to 35% of mobile phase B in 5 min, and then returned to the initial condition. Selected reaction monitoring in both positive and negative modes (spray voltage: 3200 V (pos) or 2500 V (neg); sheath gas: 35 Arb; Auxiliary gas: 20 Arb; Ion transfer tube temperature: 350 °C; vaporizer temperature: 380 °C) was used to detect the targets: dATP (492→136, pos), dGTP (508→152, pos), dCTP (466→ 158.9, neg), TTP (481→158.9, neg), ATP (508→136, pos), GTP (524→152, pos), CTP (482→158.9, neg), UTP (483→ 158.9, neg). Extracted nucleotide samples were reconstituted in 100 μL of mobile phase A. After centrifuging at 13,800 x g for 10 min, 40 μL of supernatant was mixed with 10 μL of 13C and 15N labeled dNTPs and rNTPs as internal standards, and then subjected to analysis. Data were collected and processed by Thermo Xcalibur 3.0 software. Calibration curves were generated from nucleotide standards by serial dilutions in mobile phase A (dATP and dGTP 0.1 – 400 nM, dCTP and TTP 0.2 – 400 nM, rNTPs 1 – 4000 nM). The calibration curves had r2 value greater than 0.99. All the chemicals and standards are analytical grade or higher and were obtained commercially from Sigma Aldrich (St. Louise, MO). Nucleotide standards were at least 98% pure.

### Preparation of whole cells extracts

*C. reinhardtii* cells were grown at room temperature for 3-4 days in the light in 20 mL of TAP media (ThermoFisher). The algae were collected by centrifugation at 10,000 x g for 1 min. The supernatant was removed, and the algae were washed with water and centrifuged again. The algae were resuspended in 400 μL of Algae Extraction buffer (60mM Na2CO3, 60mM DTT, 2% SDS, 12% sucrose) containing 1X cOmplete Mini, EDTA-free protease inhibitor cocktail (Roche, Milwaukee, WI). 100 μL of acid washed glass beads (425-600nm diameter) was added and then vortexed at 4 °C for 20 min. The supernatant was collected after centrifugation at 10,000 x g for 20 min at 4 °C. Aliquots were precipitated by the addition of 5 volumes of ice-cold acetone and incubated at −20 °C for 2 h. The precipitate was collected by centrifugation at 10,000 x g for 30 min at 4 °C and then air dried on ice. The dried proteins were resuspended in the appropriate buffer for further analysis. Coomassie stained gel of a 10% denaturing polyacrylamide gel was run at 120 volts for 90 min of the unpurified extraction and the purified extraction (see **Figure S1A**). Bradford assay was performed on the purified sample indicating 0.9ug/ul of protein present. SDS in the unpurified sample prevented an accurate measurement of protein concentration.

### Assays to test cell extract activity to cleave at rNMPs in DNA

Mixture of 2.5 pmol of Cy5 5’ labeled oligonucleotide containing an RNMP (Cy5.3PS.rG or Cy5.3PS.rA) or the control DNA oligonucleotide (Cy5.3PS) and 3.75 pmol of complementary oligonucleotide (DNA.comp.3PS) in 1X Thermopol Reaction Buffer was heat denatured in boiling water for 5 min and cool down at room temperature to 30 °C to anneal the complementary oligonucleotides. 2.5 pmol of annealed oligonucleotide were incubated with 400 ng of *C. reinhardtii* protein extract, or just water as negative control, for 4 hours at room temperature. Successively, the mixture was treated with 9.95 ul of formamide (VWR, 0606-100ML) and incubated for 5 min at 95 °C to denature the double strand substrate, then the mixture was put on ice. The denatured substrate was mixed with 2.22 ul of 10X Orange Loading Dye (LI-COR, C80809-01) and loaded on a 15% 7M urea denaturing polyacrylamide gel. As a positive control for this experiment, we used 5 units of *Escherichia coli* RNase HII (NEB, M0288L) in place of the *C. reinhardtii* protein extract. To test whether there was any inhibitory factor for RNase H2 activity present in the *C. reinhardtii* protein extract, we treated the 2.5 pmol of annealed oligonucleotide containing the rG with 5 units of *E. coli* RNase HII in presence or absence of 400 ng of the *C. reinhardtii* protein extract.

### Algal Genomic DNA extraction and preparation

After 5-7 days of growth in the presence of light, at a concentration of 2×10^6^ cells/mL, total DNA was isolated from 150 mL of algal cells grown in TAP medium as described in (Newman et al., 1990), with some modifications. Typically, algal cells were spun down at 3000 rpm corresponding to 1,865 x g, in three individual 50 mL sterile tubes and the pellets were resuspended in 500 μL of dH2O and combined in one sterile tube (50 mL), and 3.0 mL of SDS elution buffer (SDS 2%, NaCl 400 mM, EDTA 40 mM, Tris-HCl 100 mM, adjust pH 8.0) was added. Algal genomic DNA was extracted three times using a phenol/chloroform/isoamyl alcohol mixture (25:24:1) to eliminate all protein residues. Finally, the aqueous layer was extracted with chloroform: isoamyl alcohol (24:1). Algal genomic DNA was precipitated with isopropanol and washed with 70% cold ethanol. The pellet was air-dried overnight and dissolved in 200 μL of RNase/DNase free water. Genomic DNA was stored at −20 °C for the construction of the ribose-seq libraries.

### Ribose-seq library construction to map rNMPs in *C. reinhardtii* genomic DNA

The ribose-seq libraries were prepared as previously described with some modifications (Balachander et al., 2020; Balachander et al., 2019). Extracted algal gDNA was fragmented using three different sets of restriction enzymes (SRE) all obtained from NEB: SREI (HaeIII, HincII, and PvuII), SREII (AfeI, AluI, and SspI), and SREIII (NaeI, PsiI, and RsaI). Each set of mixtures was incubated overnight at 37 °C to produce fragments of 500- to 3,000-bp gDNA with an average size of ~1.5 kb. Then, the fragments were tailed with dATP (Sigma-Aldrich) by Klenow fragment (3’→5’ exo-) (NEB, Ipswich, MA) for 30 min at 37 °C. The resulting gDNA fragments were purified by QIAquick PCR Purification Kit (Qiagen) and then ligated to pre-annealed double-stranded adaptors that contain single dT overhangs and a unique molecular identifier (UMI) consisting of a randomized 8-base sequence containing a 3-base specific barcode by T4 DNA ligase (NEB) overnight at 15 °C. The resulting gDNA products were purified using RNAClean XP (Beckman Coulter). The adaptor-ligated DNA fragments were incubated in 0.3 M NaOH for 2 h at 55 °C to expose 2’,3’-cyclic phosphate and 2’-phosphate termini of DNA at rNMP sites, followed by neutralization and purification by using RNAClean XP. The resulting single-stranded (ssDNA) products were incubated with AtRNL buffer (50 mM Tris-HCl, pH 7.5, 40 mM NaCl, 5 mM MgCl_2_, 1 mM DTT, 30 μM ATP (Sigma-Aldrich) and 1 μM AtRNL for 1 h at 30 °C, followed by purification using RNAClean XP. The resulting products were treated with T5 exonuclease (NEB) for 1.5 h at 37 °C to degrade the background of unligated, linear ssDNA, leaving self-ligated ssDNA circles intact. Then, the purification was performed by using RNAClean XP. Samples were successively treated with 1 μM Tpt1, in Tpt buffer solution (20 mM Tris-HCl, pH 7.5, 5 mM MgCl_2_, 0.1 mM DTT, 0.4% Triton X-100), and 10 mM NAD^+^ (Sigma-Aldrich) for 1 h at 30 °C to remove the 2’-phosphate remaining at the ligation junction. After purification using RNAClean XP, the circular fragments were PCR-amplified using two rounds of amplifications to result in a ribose-seq library. A first round of PCR begins with an initial denaturation at 98 °C for 30 sec. Then denaturation at 98 °C for 10 sec, primer annealing at 65 °C for 30 sec, and DNA extension at 72 °C for 30 sec are performed; these 3 steps are repeated for 6 or 11 cycles. The 1st PCR round was performed to amplify and introduce the sequences of Illumina TruSeq CD Index primers. A second round of PCR begins with an initial denaturation at 98 °C for 30 sec. Then denaturation at 98 °C for 10 sec, primer annealing at 65 °C for 30 sec, and DNA extension at 72 °C for 30 sec are performed; these 3 steps are repeated for 9 or 11 cycles depending on the concentration of the circular ssDNAs containing the rNMPs. Successively, there is a final extension reaction at 72 °C for 2 min for both PCRs.

The primers (PCR.1 and PCR.2) (**Table S1**) used for the first PCR round were the same for all libraries. A second round of PCR was performed to attach specific indexes i7 and i5 for each library. The sequences of PCR primers and indexes can be found in **Table S1**. PCR round 1 and 2 were performed using Q5-High Fidelity polymerase (NEB) for 6 (FS231 and FS232) or 11 (FS121), and 9 (FS121) or 11 (FS231 and FS232) cycles, respectively. Following the PCR cycles, each ribose-seq library was loaded on a 6% non-denaturing polyacrylamide gel and stained using 1X SYBR Gold (Life Technologies) for 40 min.

The ribose-seq product from several PCR reactions was selectively extracted to recover 200-700 bp to exclude any primer dimers and long products that are not proficient for sequencing. The DNA was recovered from the PAGE gel using the crush and soak method (Chen and Ruffner, 1996). The resulting ribose-seq libraries were mixed at equimolar concentrations and normalized to 1.5 nM. The libraries were sequenced on an Illumina NextSeq in the Molecular Evolution Core Facility at Georgia Institute of Technology.

#### Processing and alignment of sequencing reads

The sequencing reads consist of an eight-nucleotide UMI, a three-nucleotide molecular barcode, the tagged nucleotide (the nucleotide tagged during ribose-seq from which the position of the rNMP is determined), and the sequence directly downstream from the tagged nucleotide. The UMI corresponds to sequencing cycles 1-6 and 10-11, the molecular barcode corresponds to cycles 7-9, the tagged nucleotide corresponds to cycle 12, and the tagged nucleotide’s downstream sequence corresponds to cycles 13+ of the raw FASTQ sequences. The rNMP is the reverse complement of the tagged nucleotide. Before aligning the sequencing reads to the reference genome, the reads were trimmed based on sequencing quality and custom ribose-seq adaptor sequence using cutadapt 1.16 (-q 15 - m 62 -a ‘AGTTGCGACACGGATCTATCA’). In addition, to ensure accurate alignment to the reference genome, reads containing fewer than 50 bases of genomic DNA (those bases located downstream from the tagged nucleotide) after trimming were discarded. Following quality control, the Alignment and Coordinate Modules of the Ribose-Map toolkit were used to process and analyze the reads (Gombolay et al., 2019). The Alignment Module de-multiplexed the trimmed reads by the appropriate molecular barcode, aligned the reads to the reference genome using Bowtie 2, and de-duplicated the aligned reads using UMI-tools. Based on the alignment results, the Coordinate Module filtered the reads to retain only those with a mapping quality score of at least 30 (probability of misalignment <0.001) and calculated the chromosomal coordinates and per-nucleotide counts of rNMPs. All ribose-seq libraries were then checked for background noise of restriction enzyme reads. We counted the number of reads ending with a restriction enzyme cut site, which is expected not to be generated by rNMP incorporation. Some reads captured the dAMP, which is added by dA-tailing at the restriction cut site. We summed up such background reads and calculated the percentage of background noise. All libraries had very low background (0.08% - 2.50%). To allow comparison between sequencing libraries of different read depth, the per-nucleotide coverage was calculated by normalizing raw rNMP counts to counts per hundred. The FASTQ files used as input into the Ribose-Map toolkit are available upon request.

#### Nucleotide sequence context of embedded rNMPs

Using the Sequence Module of Ribose-Map, the frequencies of the nucleotides at rNMP sites and 100 nucleotides upstream and downstream from those sites were calculated for the mitochondrial, chloroplast and nuclear genomes of *C. reinhardtii*. The Sequence Module normalizes the nucleotide frequencies to the frequencies of the corresponding reference genome.

#### Heatmaps

To generate the mononucleotide heatmaps for every mitochondrial, chloroplast and nuclear ribose-seq library, the number of each type of rNMP (*R_N_*: *R_A_*, *R_C_*, *R_G_* or *R_U_*) was counted and divided by the

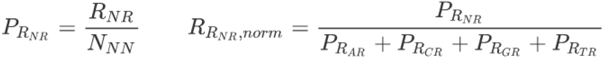

total number of rNMPs to yield the proportion *R_N_R_,raw_*:

Then, each raw counts of rNMPs (*R_N_*) was divided by the corresponding deoxy-mononucleotide frequency of the reference genome (*N_N_*) to yield the probability of rNMP incorporation (*P_R_N__*), and were normalized to

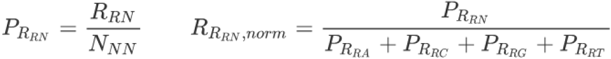

generate the normalized proportion *R_R_N_,norm_*. These data were used in the normalized mononucleotide heatmaps:

Similarly, to generate the normalized dinucleotide heatmaps, each raw count of dinucleotides with an rNMP along with a deoxyribonucleotide at position −1, −2, −3, −4, −5, −6 or −100 relative to the rNMP (*R_NR_*), or at position +1, +2, +3, +4, +5, +6 or +100 relative to the rNMP (*R_RN_*) were divided by the corresponding deoxy-dinucleotide frequency of the reference genome (*N_NN_*) to yield the probability of NR and RN dinucleotide incorporation (*P_R_NR__*) and *P_R_RN__*, respectively. Next, these proportions were normalized keeping fixed the deoxyribonucleotide base in the position of the rNMP to generate the normalized proportion

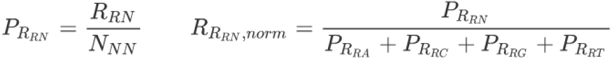

*R_R_NR_,norm_* or *R_R_RN_,norm_*

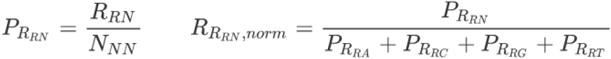

The specific background frequencies of mononucleotides and dinucleotides for mitochondrial, chloroplast and nuclear DNA are shown in the heatmap figures.

## QUANTIFICATION AND STATISTICAL ANALYSIS

### Statistical analysis for heatmaps

To compare NTP/dNTP ratios shown in Figure 1B, we used the t-test. The Chi-square test is used for analyses of heatmap data to check if the rNMPs are incorporated randomly, as described in the legend of Table S3.

## Supplementary Table Legends

**Table S1.**
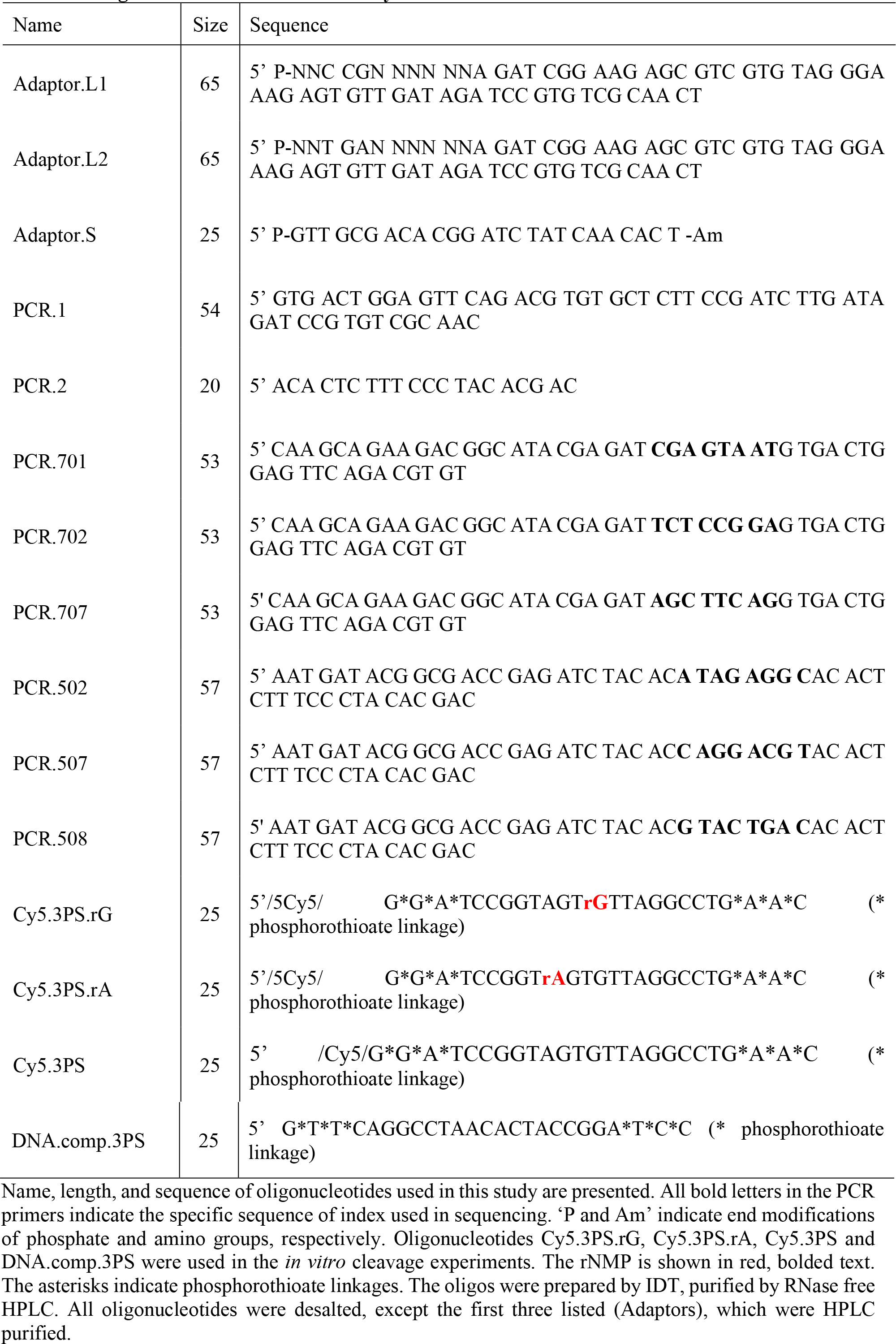
Oligonucleotides used in this study. Name, length, and sequence of oligonucleotides used in this study are presented. All bold letters in the PCR primers indicate the specific sequence of index used in sequencing. ‘P and Am’ indicate end modifications of phosphate and amino groups, respectively. Oligonucleotides Cy5.3PS.rG, Cy5.3PS.rA, Cy5.3PS and DNA.comp.3PS were used in the *in vitro* cleavage experiments. The rNMP is shown in red, bolded text. The asterisks indicate phosphorothioate linkages. The oligos were prepared by IDT, purified by RNase free HPLC. All oligonucleotides were desalted, except the first three listed (Adaptors), which were HPLC purified.

**Table S2. rNMPs found in mitochondrial, chloroplast and nuclear ribose-seq libraries of *C. reinhardtii*.** List of mitochondrial, chloroplast and nuclear ribose-seq libraries of *C. reinhardtii* constructed in this study with corresponding data information and indication of algal strain name, genotype, library name, restriction enzyme set used, number of rNMPs, % rA, rC rG and rU with mean and standard deviation for each cell compartment, barcode, and number of cycles in PCR 1 and PCR 2.

**Table S3. Comparison of ratios with *P* values of heatmap data.** Chi-square test is used for (**A**) mitochondrial, (**B**) chloroplast, and (**C**) nuclear DNA to check if the rNMPs are incorporated randomly. The observation value is the count of incorporated rNMP (mono) or dinucleotide with an rNMP (NR or RN, ±1 - ±6, ±100). The expectation is calculated with the corresponding dNMP or dNMP pair background frequency with the same number of total rNMPs. A small P-value (<0.05) means the observation is significantly different with the expectation, which suggests rNMP incorporation is not random. Due to the limit of Excel precision, a P-value less than 10^-307^ is shown as 0.

## Supplementary Figure Legends

**Figure S1. Protein extract from *C. reinhardtii* do not cleave at rNMPs in DNA.** (**A**) Coomassie stained gel of a 10% denaturing polyacrylamide gel was run at 120 volts for 90 min of the unpurified extraction and the purified extraction. The first lane contains 10ul of unpurified protein in extraction buffer containing 1X cOmplete Mini, EDTA-free protease inhibitor cocktail (Roche) while the second lane is 10ul of purified extract via acetone precipitation. (**B**) Double-stranded Cy5-labeled 25-mers containing an rNMP (rG or rA) were used in the *in vitro* cleavage assay to test RNase H2 activity *in C. reinhardtii* protein extracts, schemes shown to the left. The red arrow shows the cleavage position by RNase HII/2. N, negative control with the double-stranded oligonucleotide (with rG or rA, as indicated) treated by water. HII, *Escherichia coli* RNase HII was used as a positive control cleaving 5’ of the rNMP embedded in the double stranded DNA oligonucleotides. P, *C. reinhardtii* protein extract. (**C**) *E. coli* RNase HII (HII) and *C. reinhardtii* protein extract (P) were combined (HII + P) to test any inhibitory effect of P on RNase H activity on the rG-containing substrate (shown on the left). DNA, negative control double-stranded, DNA-only, Cy5-labeled 25-mer (shown on the left). L, ladder with 25mer and 12mer bands. N, negative control with the double-stranded oligonucleotide with rG treated by water. The cleavage % is shown underneath the image.

**Figure S2. Like the +1, the deoxyribonucleotide at positions −2 to −6, and +2 to +6 have less impact than the one immediately upstream (−1) on rAMP occurrence in *C. reinhardtii* mitochondrial and chloroplast DNA.** Heatmap analyses with normalized frequency of N-R, R-N, N--R, R--N, N---R, R---N, N----R, R----N, N-----R, R-----N, N-99-R, and R-99-N dinucleotides (rA with the −2, +2, −3, +3, −4, +4, −5, +5, −6, +6, −100 or +100 deoxyribonucleotide with base A, C, G or T) for all the mitochondrial (**A**) and chloroplast (**B**) ribose-seq libraries of this study. The formulas used to calculate these normalized frequencies are shown and explained in Materials and Methods. Each column of the heatmaps shows results of a specific ribose-seq library. Each library name is indicated underneath each column of the heatmaps. Each row shows results obtained for a dinucleotide RN, N-R, R-N, N--R, R--N, N---R, R---N, N----R, R----N, N-----R, R-----N, N-99-R, or R-99-N of fixed rNMP base A for each library. The actual % of dinucleotides of fixed base A for the indicated base combinations that are present in mitochondrial (A) or chloroplast (B) DNA of *C. reinhardtii* are shown to the left of the heatmaps. The observed % of dinucleotides with rNMPs with base A were divided by the actual % of each dinucleotide with fixed base A in mitochondrial or chloroplast DNA of *C. reinhardtii*. The bar to the right shows how different frequency values are represented as different colors. Black corresponds to 0.25.

**Figure S3. Like the +1, the deoxyribonucleotide immediately downstream (+1) from each rNMP and those at positions −2 to −6, and +2 to +6 have less impact than the one immediately upstream (−1) on rNMP occurrence in *C. reinhardtii* nuclear DNA.** Heatmap analyses with normalized frequency of N-R, R-N, N--R, R--N, N---R, R---N, N----R, R----N, N-----R, R-----N, N-99-R, and R-99-N dinucleotides (rA, rC, rG, or rU with the −2, +2, −3, +3, −4, +4, −5, +5, −6, +6, −100 or +100 deoxyribonucleotide with base A, C, G or T) for all the nuclear ribose-seq libraries of this study. The formulas used to calculate these normalized frequencies are shown and explained in Materials and Methods. Each column of the heatmaps shows results of a specific ribose-seq library. Each library name is indicated underneath each column of the heatmaps. Each row shows results obtained for a dinucleotide RN, N-R, R-N, N--R, R--N, N---R, R---N, N----R, R----N, N-----R, R-----N, N-99-R, or R-99-N of fixed rNMP base A, C, G or U for each library. The actual % of dinucleotides of fixed base A, C, G or T for the indicated base combinations that are present in nuclear DNA of *C. reinhardtii* are shown to the left of the heatmaps. The observed % of dinucleotides with NMPs with base A, C, G or U were divided by the actual % of each dinucleotide with fixed base A, C, G or T in nuclear DNA of *C. reinhardtii*. The bar to the right shows how different frequency values are represented as different colors. Black corresponds to 0.25.

