## Supplementary material for "Disproportionate presence of adenosine in mitochondrial and chloroplast DNA of *Chlamydomonas reinhardtii*": Suppl Figures S1 to S3

Figure S1

A

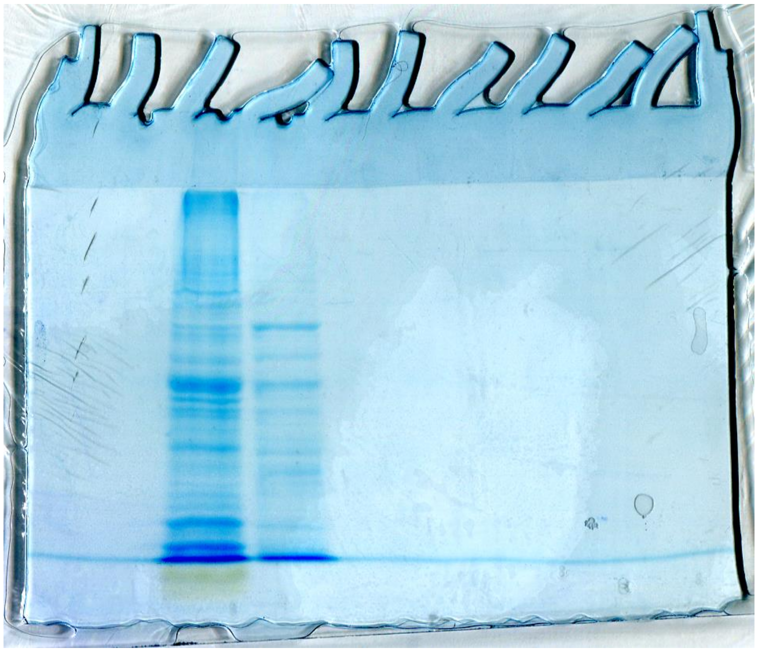

B

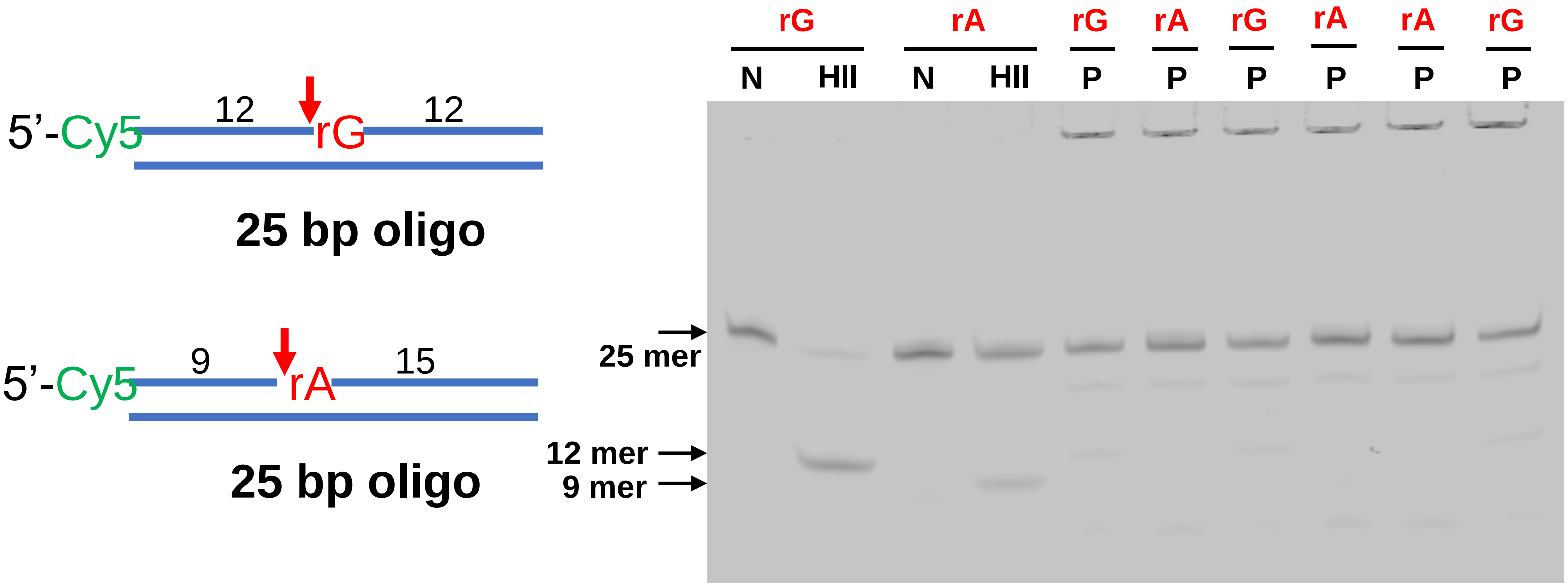


C

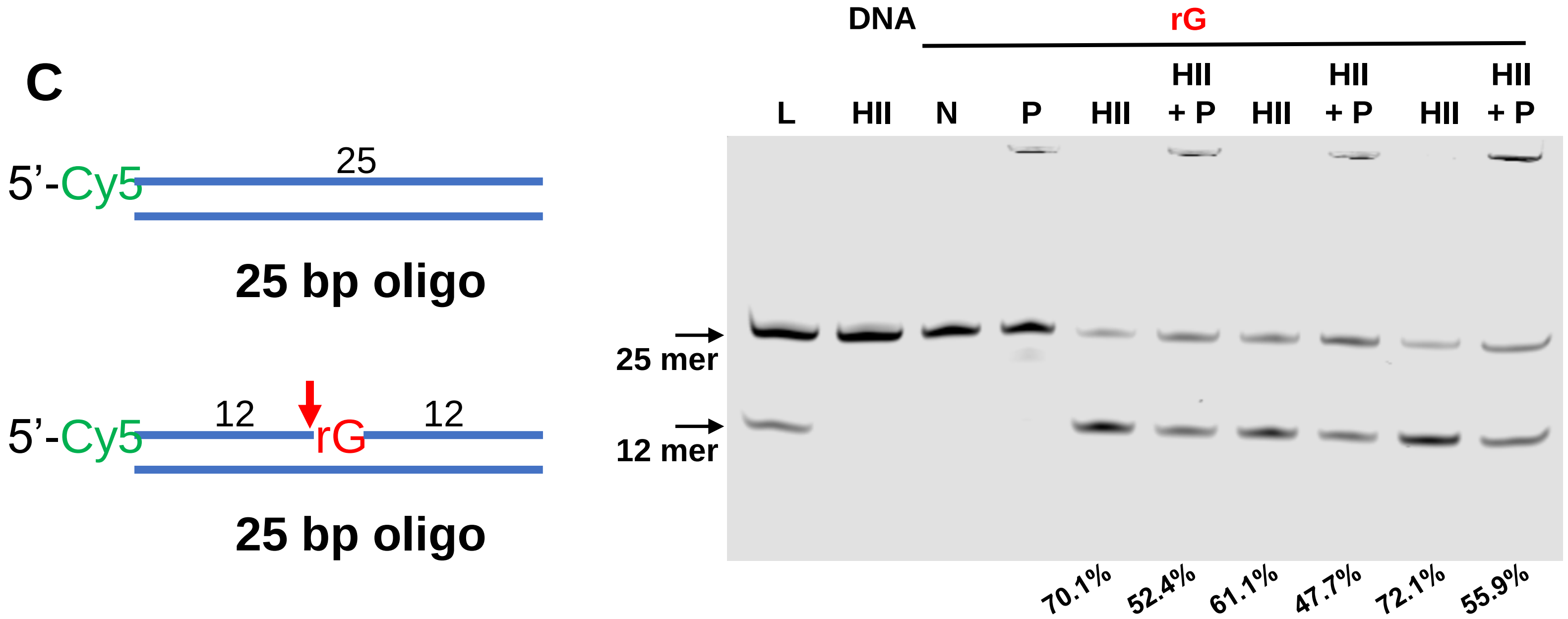

### Figure S2

## A

##### Dinucleotide ratios normalized to background frequency in mitochondria

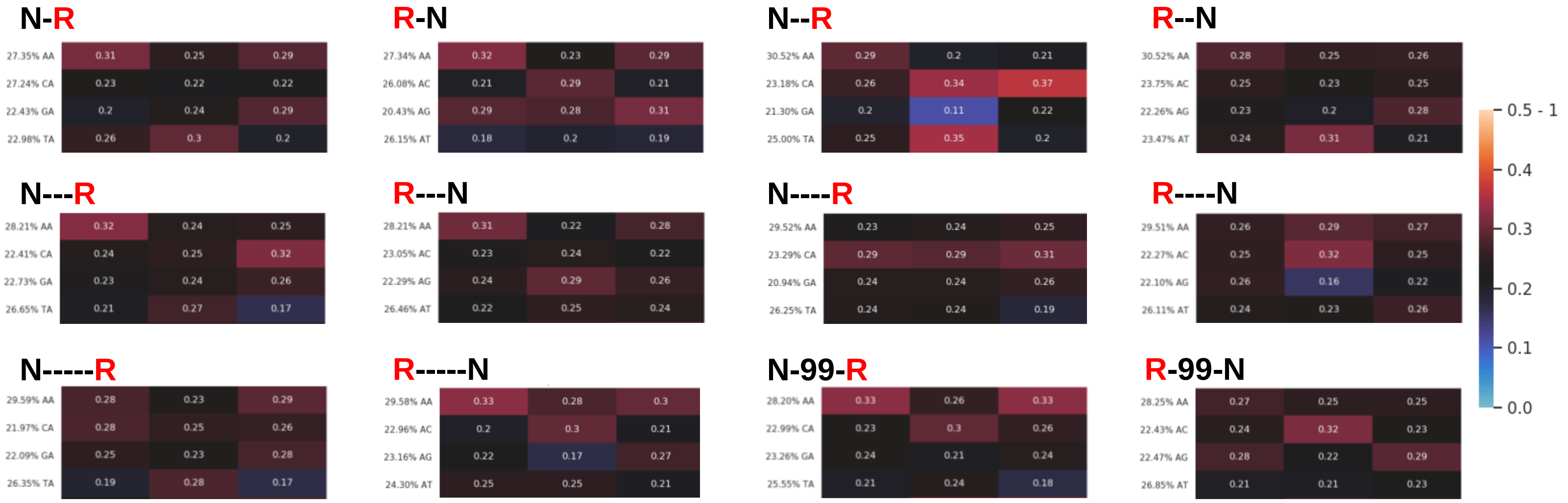


## B

##### Dinucleotide ratios normalized to background frequency in chloroplast

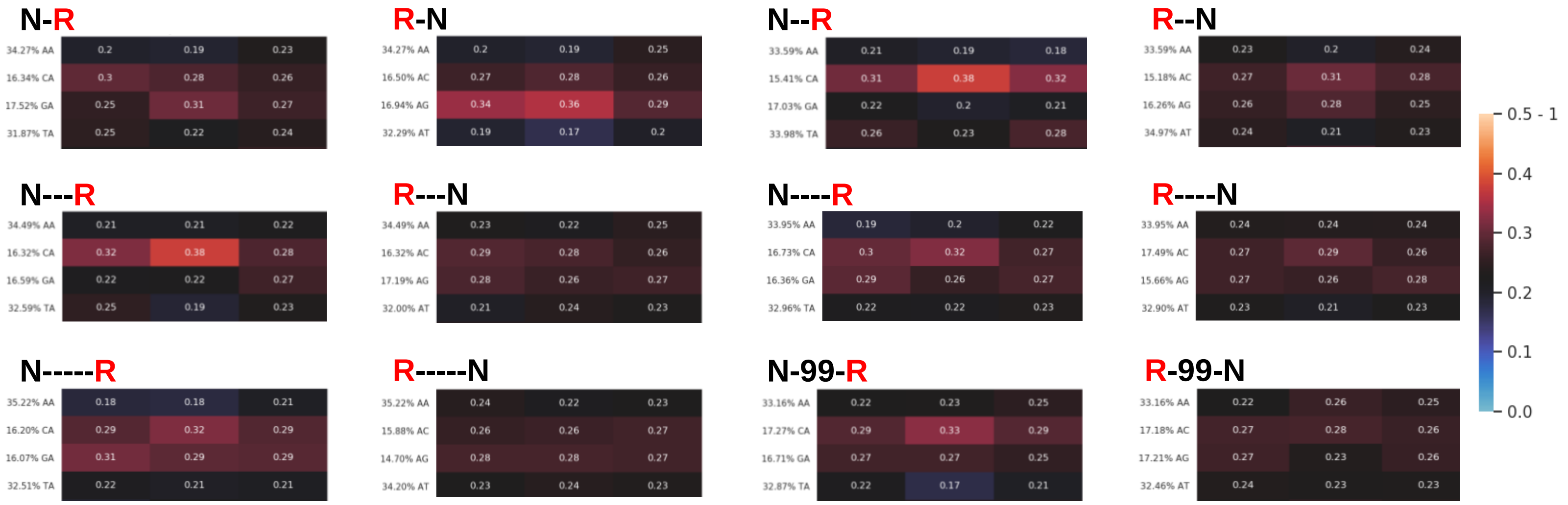

### Figure S3

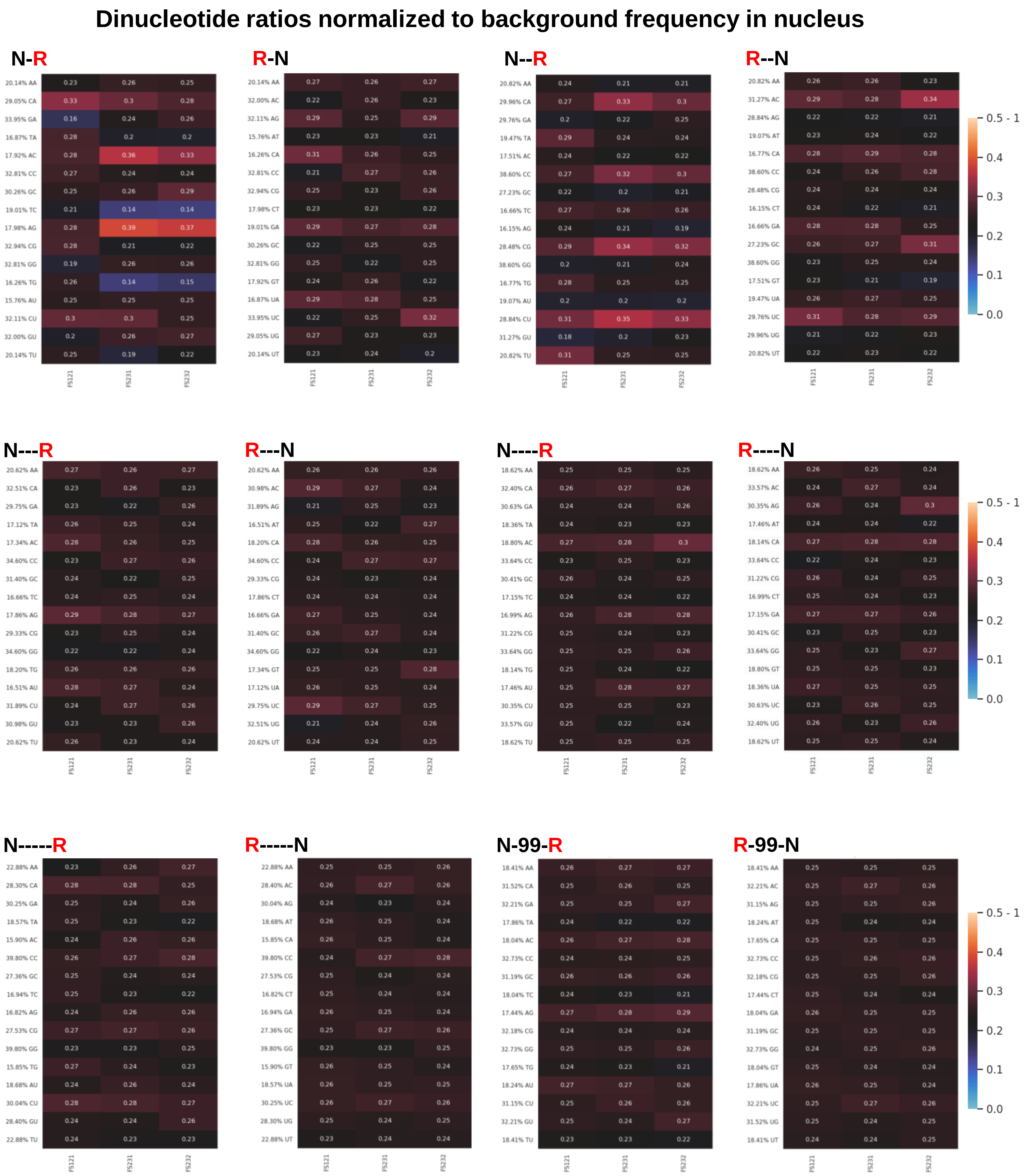
Dinucleotide ratios normalized to background frequency in nucleus
N-R
R-N
N--R
R--N
N---R
R---N
N----R
R----N
N-----R
R-----N
N-99-R
R-99-N
FS121
FS231
FS232
0.5 - 1
0.4
0.3
0.2
0.1
0.0
